# Bi-compartmentalized stem cell organization of the corneal limbal niche

**DOI:** 10.1101/2020.06.25.171462

**Authors:** Olivia Farrelly, Yoko Suzuki-Horiuchi, Megan Brewster, Paola Kuri, Sixia Huang, Gabriella Rice, Jianming Xu, Tzvete Dentchev, Vivian Lee, Panteleimon Rompolas

## Abstract

Stem cells exist in precise locations within tissues, yet how their organization supports tissue architecture and function is poorly understood. The limbus is the presumptive stem cell niche of the corneal epithelium. Here, we visualize the live limbus and track the activity of single stem cells in their native environment by 2-photon microscopy. We identify previously unknown niche compartments and show that long implicated slow-cycling cells form separate lineages in the outer limbus, with only local clonal dynamics. Instead, we find distinct stem cells in the pericorneal limbus to be required for corneal regeneration. Unbiased photolabeling captures their progeny exiting the niche, then moving centripetally in unison before undergoing terminal differentiation. This study demonstrates how a compartmentalized stem cell organization coordinates tissue regeneration.

**One Sentence Summary:** In vivo live imaging of the regenerating cornea reveals distinct stem cell activities in the limbal niche

---

Stratified squamous epithelia provide a barrier for organs that are directly exposed to the environment. To support their function, these tissues undergo continuous regeneration, characterized by high cellular turnover. During this process, terminally differentiated cells are shed from their surface and are replenished by resident stem cells located in the basal layer (*1, 2*) In most stratified squamous epithelia, including the epidermis and esophagus, stem cells lack a distinctive niche and are instead distributed throughout the tissue (*3, 5*). In contrast, the cornea has a spatially defined niche at its periphery, known as the limbus (*6, 8*) The cornea, which lines the anterior portion of the eye, is compartmentalized from the rest of the surface epithelium and is characterized by a unique physiology (*9*). While stem cells in the limbus are critical for corneal function and maintaining vision, their organization and distinct behaviors within their native niche are poorly understood.

Resolving the cellular dynamics of stem cells within their niche is critical to understand how tissues can regenerate while maintaining functional organization (*10*). Experimental and clinical evidence suggest that the limbus harbors bona fide stem cells that regenerate the cornea, and is also the location of slow-cycling, label-retaining cells, long presumed to be part of the same corneal stem cell hierarchy (*11-17*). Slow-cycling ability is a hallmark of stem cells in various organs, but is also associated with distinct reserve populations (*18-21*). The function of slow-cycling cells in the limbus and how they contribute to tissue regeneration have not been elucidated. A significant challenge to resolving the cellular mechanisms of corneal regeneration has been the absence of means to directly visualize, track and test the activity of individual stem cells in their native environment. To overcome this limitation, we established a technique to non-invasively image the eyes of live mice by 2-photon microscopy and capture dynamic behaviors in the limbal niche at single-stem-cell resolution (*22-24*) **(Fig. 1A, Fig. S1A; Movie S1)**.

**Fig. 1.**
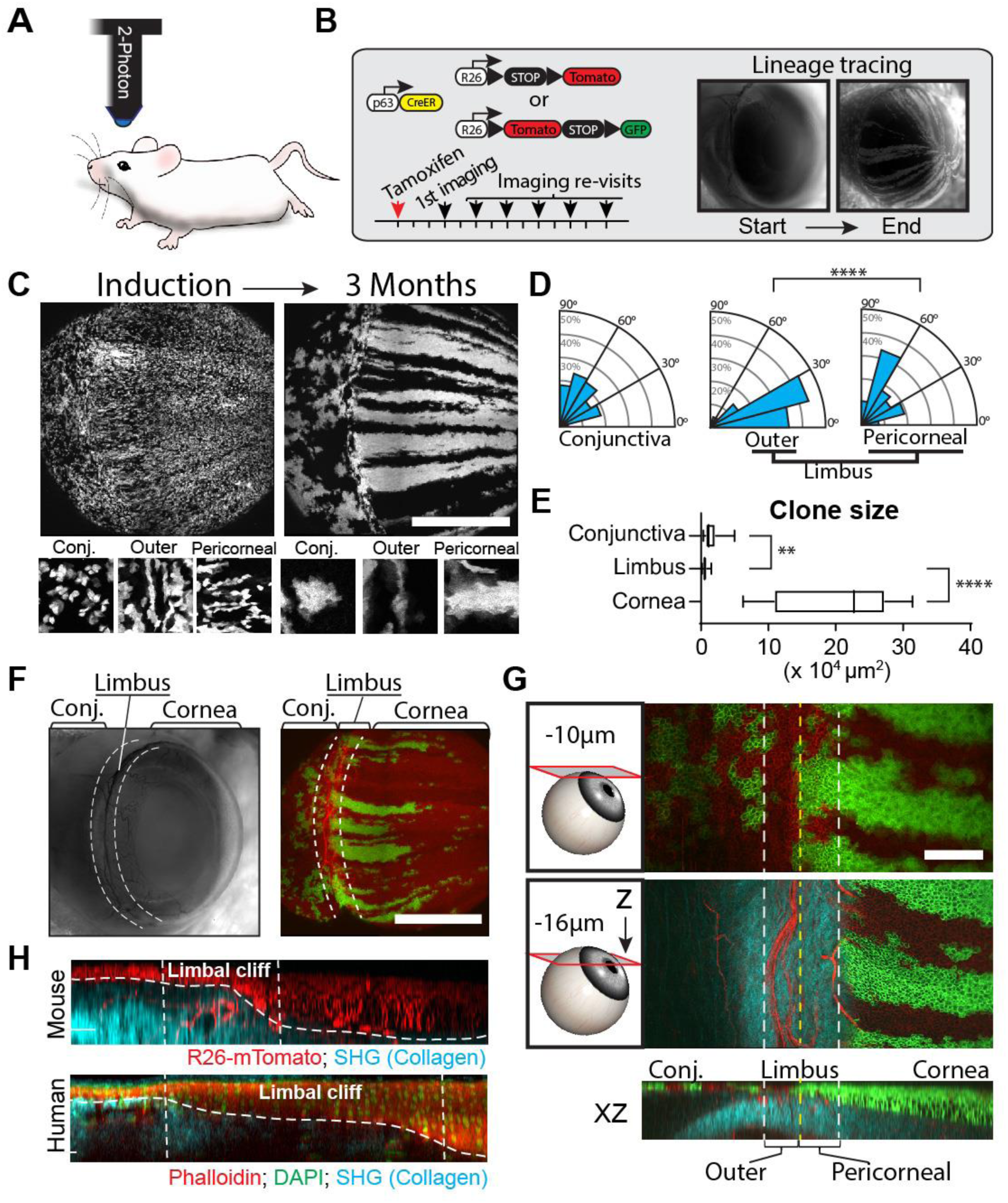
Intravital imaging of the limbal niche reveals distinct clonal growth patterns. **(A)** Single-cell resolution imaging of eyes in live mice is performed with 2-photon microscopy. **(B)** Experimental strategy to resolve clonal dynamics in the eye surface epithelium by live imaging and lineage tracing. **(C)** Global and high-magnification views of the eye surface epithelium imaged at the indicated time points. Individual stem cells and basal progenitors (p63+) are marked uniformly in all epithelial compartments at the time of induction. After three months, unique clonal patterns emerge in the cornea, limbus and conjunctiva. **(D)** Quantification of clonal growth anisotropy, five days after induction, is depicted in the radial graphs (N = 206; \*\*\*\**P < 0*.*0001*). **(E)** Quantification of clone size, three months after induction (N = 45; \*\**P* = 0.0079, \*\*\*\**P < 0*.*0001*). **(F)** Low magnification side view of the anterior mouse eye segment imaged live by wide-field (left) or 2-photon microscopy (right). **(G)** High magnification views of the intact live limbus. Two representative optical planes at the indicated depth from the surface and a reconstructed side view (XZ) are shown to illustrate the unique anatomical characteristics and clonal organization of the niche. The outlines of all cells are visualized in red (Tomato) and collagen fibers of the extracellular matrix in blue (SHG). Long-lived epithelial clones are shown in green (GFP). **(H)** Side views (XZ) of the mouse and human limbus, reconstructed from full-thickness serial optical sections. Scale bars: 1 mm (C, F), 200 μm (G).

Due to the lack of definitive molecular markers, the limbus has instead been recognized anatomically as the border between the cornea and conjunctiva, the two epithelial compartments of the eye that form a continuous surface (*25*). To gain access to the live limbal niche without compromising the structural integrity or physiology of the eye, we custom-built a microscope stage to mount and stabilize the anesthetized mouse during the imaging process **(Fig. S1A)**. To resolve the cellular organization of the limbus, we used mice that ubiquitously express fluorescent reporters to visualize the plasma membrane or nuclei of all cells (**Fig. S1B, C; Movie S2, Movie S3)**. In addition, we used second harmonic generation (SHG) microscopy to visualize the organization of the collagen fibers in the extracellular matrix (*26*) (**Fig. S1B; Movie S4)**. Together, these enabled the precise 3-dimensional analysis of the intact limbal niche and the direct examination of the cellular organization across the anterior surface epithelium of the eye.

The cornea consists of 8-10 discrete epithelial cell layers, but changes abruptly to only 2-3 layers in the limbus before transitioning to the conjunctiva **(Fig. S1C)**. Due to its appearance, we refer to this transition zone as the “limbal cliff”. Unique characteristics of this area include a dense network of blood vessels and nerves that extend circumferentially along the limbal cliff, and a stroma with a network of distinctly-oriented collagen fibers **(Fig. S1B, C)**. While the cell density in the basal and suprabasal layers is uniform throughout the entire cornea, it decreases dramatically in the limbus and the conjunctiva **(Fig. S1C)**.

Lineage tracing studies suggest that the cornea is replenished by long-lived clones that originate from its periphery, and is therefore presumed to be self-contained in its ability to regenerate (*15-17*). To determine the precise location of the stem cells and to resolve the clonal dynamics that enable the compartmentalization of the cornea, we devised a Cre-recombinase based lineage tracing strategy combined with recurrent live imaging **(Fig. 1B; S2A)**. Using a conditional p63 genetic driver, we uniformly marked basal stem/progenitor cells across the entire surface epithelium, comprised of the cornea, limbus and conjunctiva **(Fig. S3A)**. To track these cells and determine their fate, we first acquired high-resolution, full-thickness serial optical sections of the whole eye including the limbus, and then re-imaged the same eyes over time using identical acquisition parameters **(Fig. S2A, B)**. Several weeks after induction, only clones that were sustained by stem cells were present in the tissue (**Fig. 1C, F; S2B; S3A)**. Overall, this suggests that our labeling and imaging strategy allows us to discriminate between long-lived stem cells and their transient progeny by tracking their clonal evolution over time.

Analysis of the lineage tracing data classified clones into three major categories based on their location, dimensions, and orientation **(Fig. 1C-G; S2B-D)**. In the conjunctiva, long-lived lineages were found in differing sizes and shapes that are reminiscent of neutrally competing clones described in the epidermis and the esophagus (**Fig. 1D, E)**. This suggests that the conjunctiva follows a similar stem cell organization as has been proposed for these epithelia (*3, 5, 27*) In the limbus, we found evidence of at least two distinct stem cell activities. Large radially-oriented clones that regenerated the cornea had a common point of origin at the base of the limbal cliff **(Fig. 1G-H; S2B-D)**. This compartment, which we refer to as the “pericorneal limbus”, harbors the corneal stem cells. Smaller spatially-distinct clones formed in a narrow zone immediately adjacent in the “outer limbus” **(Fig. 1C-G; S2B-D)**. These limbal clones co-localized and orientated parallel to the circumferentially aligned blood vessels and collagen fibers of the underlying stroma **(Fig. 1E-G; S2B-D)**. Analysis of the anisotropy of the growth dynamics in early clones further supports the bi-compartmentalized organization of the limbus (**Fig. 1D; S3B)**. Importantly, the anatomical and cellular characteristics of the limbus appear to be conserved between mice and humans **(Fig. 1H; Movie S5)**.

Terminal differentiation in all squamous stratified epithelia induces a continuous flux of cells that leave the basal layer and transit upwards to replace those shed from the surface of the tissue (*3, 5, 28*). The large radial clonal patterns in the cornea point to a secondary centripetal flux of cells within the basal progenitor population. The potency and long-term contribution of basal progenitors in the cornea have been the subject of intense research (*29-30*). We next tested the hypothesis that transit amplifying progenitors in the central cornea are replenished by stem cells that reside exclusively in the pericorneal limbus. Using a globally expressed photo-activatable mouse reporter and live imaging, we selectively marked basally located cells in all epithelial compartments and tracked their activity over time **(Fig. 2A; Movie S6)**.

**Fig. 2.**
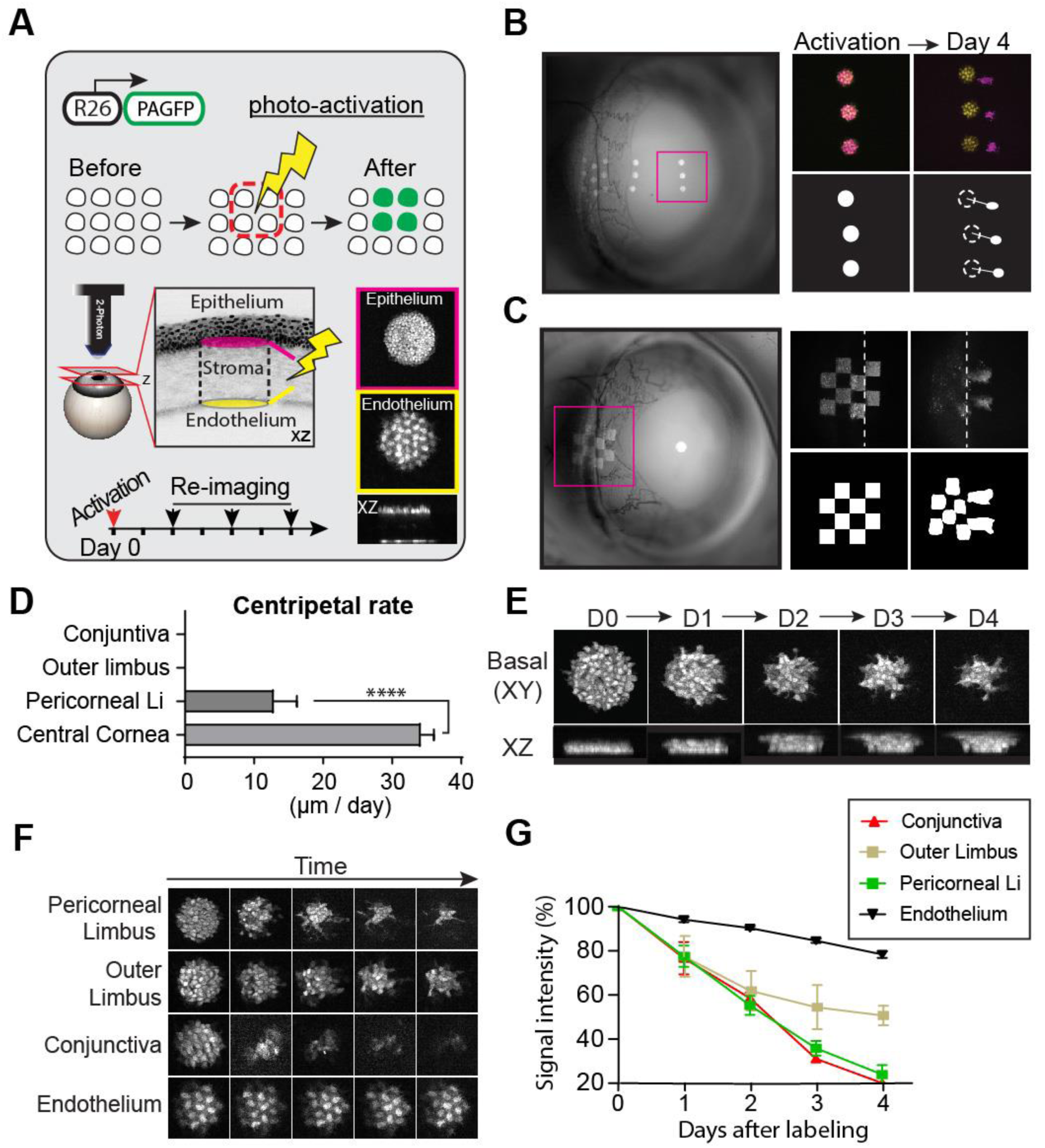
Photo-labeling and tracking of stem cell dynamics in the live eye. (**A)** Experimental strategy to directly capture the mobility and fate of stem cells and basal progenitors in the eye surface epithelium. Basal cells are marked in an unbiased manner using a genetically encoded and globally expressed, photo-activatable reporter (PAGFP). Post-mitotic, corneal endothelial cells are used as a reference. The movement and fate of marked cells are tracked over time by re-imaging the same areas. **(B)** Low magnification, wide-field fluorescent image of the eye showing the location of photo-labeled cells (left panel). Full-thickness projections of serial optical sections acquired by 2-photon microscopy. Basal corneal progenitors (magenta) co-localize with endothelial cells (yellow) immediately after photo-activation, but show uniform centripetal translocation after 4 days. Area of each photo-labeled group at “Day 0” is 100 μm in diameter. **(C)** Basal cells in the limbal area are marked in a checkered pattern to visualize the local cellular dynamics. Pattern deformation shows centripetal expansion in the pericorneal limbus (dotted line) but not the outer limbus or conjunctiva. **(D)** Quantification of cellular translocation rates across the eye epithelium (*N* = 44; \*\*\*\**P* < 0.0001). **(E)** Representative, high magnification, single optical sections (top panel) and reconstructed side views (bottom panel), showing cells initially photo-labeled at the basal layer and tracked at daily time intervals. During the time course, basal cells that undergo terminal differentiation move towards the suprabasal layers. **(F)** Panel shows representative images of photo-labeled basal cells from the indicated epithelial compartments. The same groups of cells were re-imaged at daily time intervals. A group of quiescent corneal endothelial cells (bottom row) were used as a control. **(G)** Quantification of basal cellular turnover as a function of PAGFP signal dilution. (*N* = 15; *\*\*\*\*P* < 0.0001, 2-way ANOVA)

We found that, in the cornea, all cells move unidirectionally towards the center at defined rates (**Fig. 2B, D; S4; S5, Movie S6)**. The concerted centripetal mobility of corneal progenitors leads to their terminal differentiation due to the constraints imposed by the circular geometry of the tissue. In contrast, distinctly labeled groups of cells in the pericorneal limbus remain in their original location, but also give rise to progenitors that expand centripetally **(Fig. 2C; S6)**. This indicates that stem cells in the pericorneal limbus are the source of all basal cells that populate the rest of the cornea. Taken together, our data directly demonstrate that the corneal progenitors that exit the pericorneal limbal niche are short-lived and have finite contributions to regeneration. In contrast, marked cells in the limbus and conjunctiva do not display lateral mobility or expansion **(Fig. 2C; S5)**. We used the rate of label dilution to measure cellular turnover due to proliferation and differentiation. This showed distinct dynamics in each of the epithelial compartments, consistent with the lineage tracing data **(Fig. 2E-G)**. Basal cells labeled in the outer limbus showed a slower overall turnover than those immediately adjacent in the pericorneal limbus and conjunctiva **(Fig. 2G)**. Together, these data further solidify the conclusion that cells in the pericorneal limbus are likely the definitive stem cells from which transit amplifying progenitors that regenerate the cornea ultimately emerge.

We next wanted to visualize the previously reported label-retaining cells in order to define their precise location within the limbus and resolve their role in corneal regeneration (*7, 31*). For this, we first engineered transgenic mice that express a photo-activatable GFP fused to histone-H2B under the control of a Keratin 14 promoter, which shows particularly high activity within the limbus (*32, 33*) **(Fig. 3A, Fig S7A, B)**. Photoactivating the reporter produces a robust nuclear signal that is stable and diluted equally among daughter cells after each cell division (*5, 19*). After a chase period, label-retaining GFP+ cells were present only in the outer limbus **(Fig. 3B; S7D)**. This suggests that these slow-cycling, label-retaining cells are functionally distinct from the corneal stem cells we identified in the pericorneal limbus. To further test this, we bred mice that combined the histone-H2B photo-activatable reporter with the p63 lineage tracing alleles. GFP+ cells co-localized with the small clones in the outer limbus, but were not part of the lineages that emerged from the pericorneal limbus **(Fig. 3C; S7E)**. Therefore, our data thus far support a previously unrecognized bi-compartmentalized organization of the limbal niche and show that label-retaining cells located in the outer limbus are functionally distinct from corneal stem cells **(Fig. 3D)**.

**Fig. 3.**
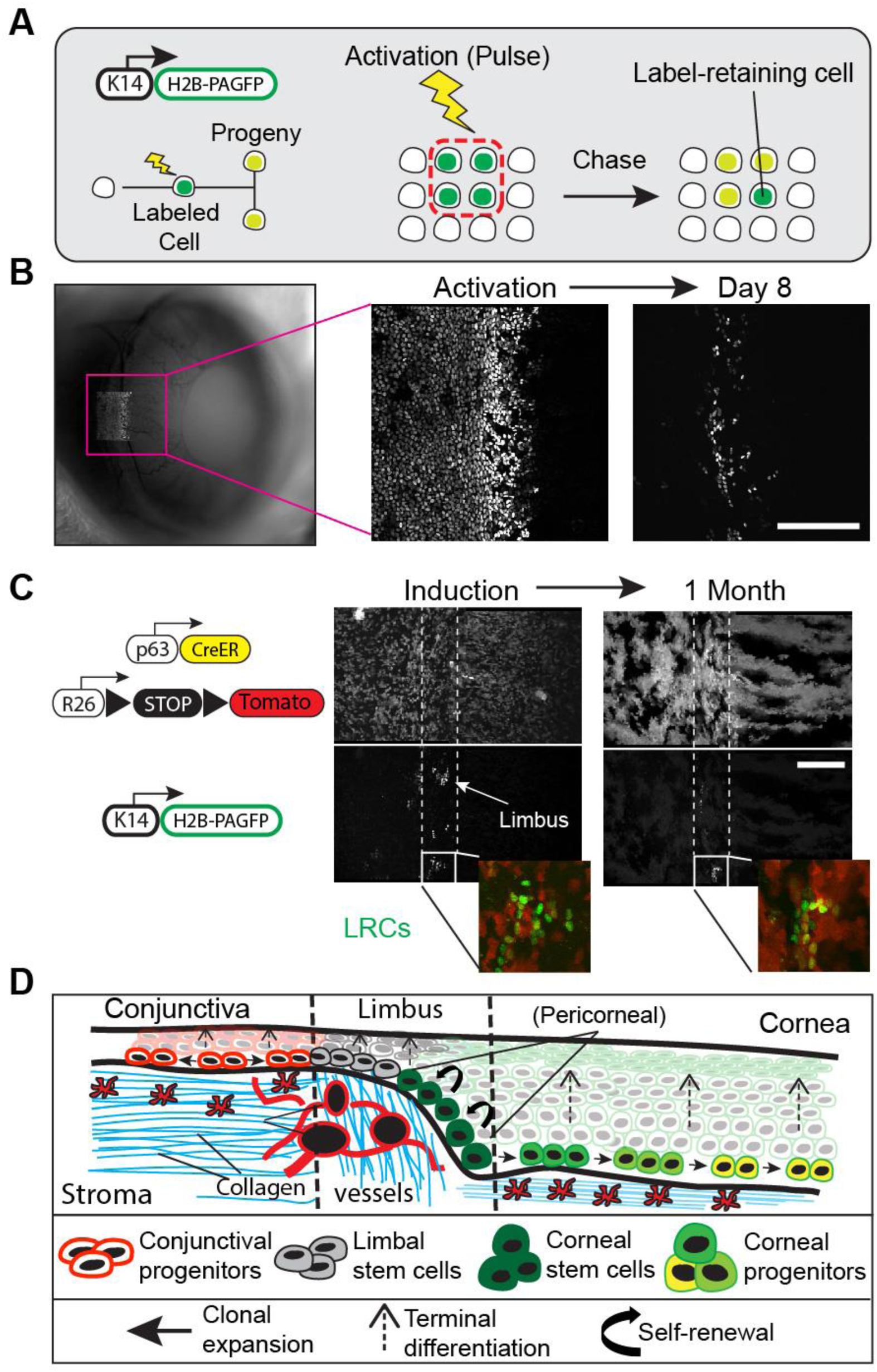
Label-retaining cells in the limbus are distinct from corneal stem cells. **(A)** Experimental strategy to visualize label-retaining cells in the eye by live imaging. H2B-PAGFP expressing cells in the limbal area are marked by photo-activation and the same cells are re-imaged after a chase period. The nuclear signal is diluted among daughter cells and only cells that have undergone the fewest cell divisions will retain their label and be visible after chase. **(B)** Low magnification wide field fluorescent image of the eye immediately after photo-labeling cells in the limbal area (left panel). High magnification views of the same photo-labeled cells at the beginning and end of the chase period, acquired by 2-photon live imaging (right panel). Label-retaining cells are localized exclusively in the outer limbus. **(C)** Representative images of the limbal area taken at the time of tamoxifen induction and after one month. H2B-PAGFP+ cells do not co-localize with stem cells in the pericorneal limbus. Dotted lines indicate the margins of the outer limbus (white) and the pericorneal limbus. **(D)** Model of the observed stem cell activities in the limbal niche and across the eye surface epithelium. Scale bars: 200 μm.

To test whether each stem cell population is required for corneal regeneration, we specifically removed cells by laser-mediated ablation and tracked clonal capacity and replacement potential by neighboring cells **(Fig. 4A)**. Using high magnification optics, epithelial cells were specifically ablated without disrupting the overall organization of the niche (*22*) **(Fig. 4B, S8A)**. Ablation of corneal progenitors away from the pericorneal limbal niche did not impede the growth of their respective corneal lineage (**Fig. S8)**. Surprisingly, the centripetal motion of the progeny downstream of the ablation site continued until those cells were eventually exhausted through terminal differentiation (**Fig. S8)**. After specific ablation of cells in the pericorneal limbus, the established large radial clones failed to sustain growth and regressed centripetally. These data directly demonstrate that stem cells in the pericorneal limbus are required to regenerate the cornea **(Fig. 4C, D)**.

**Fig. 4.**
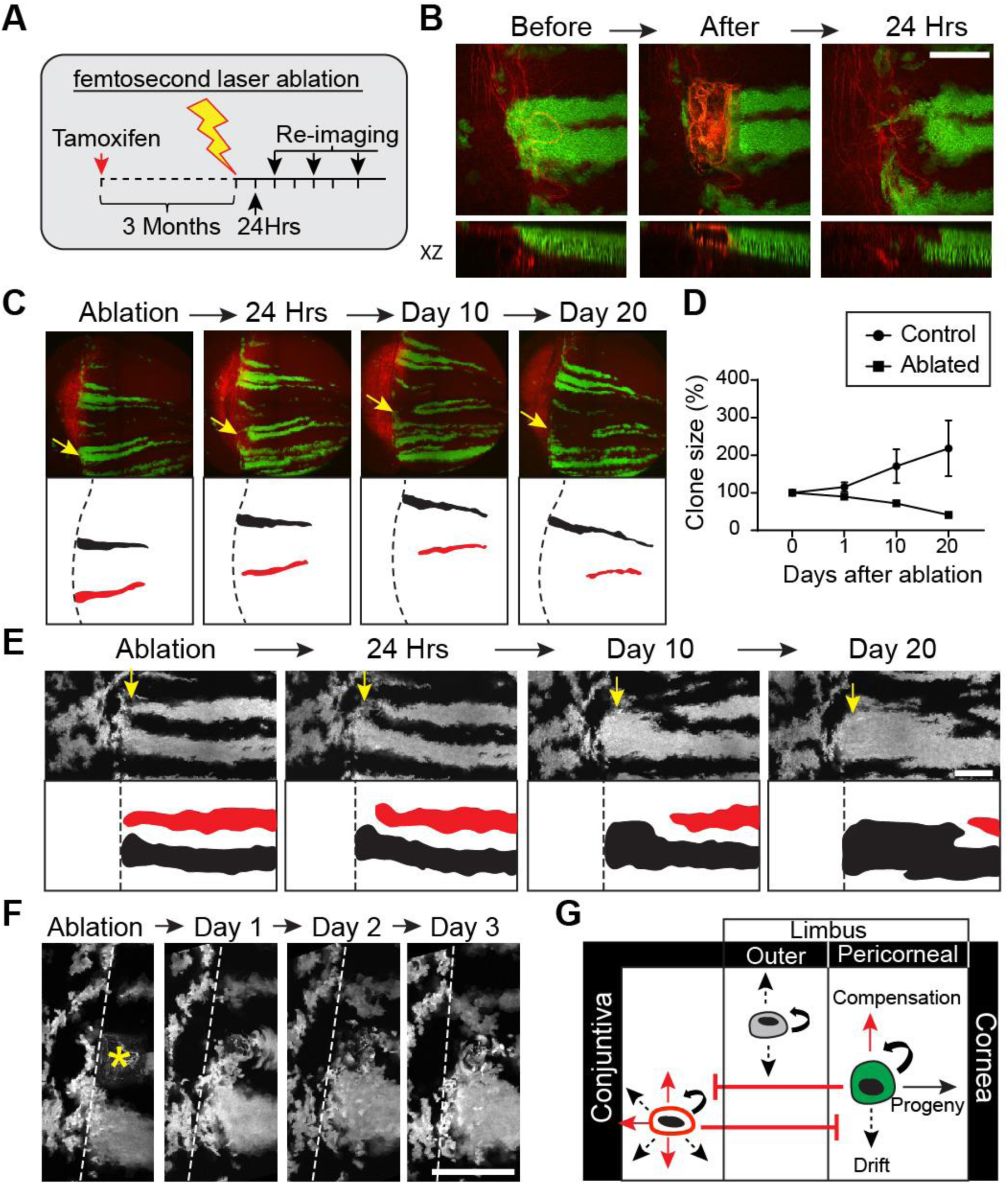
Compartmentalized stem cell requirement and contribution to regeneration. **(A)** Experimental strategy to test stem cell requirement and contribution to regeneration by live imaging. Long-lived clones are first allowed to form for at least 3 months after tamoxifen induction. A femtosecond laser is then used to specifically ablate stem cells with minimal disruption to the limbal niche. The fate of clones after stem cell ablation is determined by re-imaging the same areas at defined time intervals. (**B)** Single optical sections (top) and reconstructed side views (bottom) of the limbal area taken at the indicated time points before and after ablation. Note that the underlying limbal niche is not affected by the ablation of the corneal stem cells. **C**. Stem cells in the pericorneal limbus are required for regeneration. Low magnification views of the entire eye, imaged live by 2-photon at the indicated time points (top panel). Traces of individual corneal lineages with intact (black) or ablated (red) pericorneal stem cells (bottom panel). Arrows indicate the site of ablation and the dotted lines mark the limbal border. (**D)** Quantification of the size of corneal clones with intact or ablated pericorneal stem cells. (*N = 6; P* = 0.0062, 2-way ANOVA) (**E)** Stem cells from proximal corneal clones compensate for cell loss after ablation. Representative views of the limbal area taken at the indicated timepoints after stem cell ablation. A corneal lineage (red trace) regresses after ablation of its associated stem cells. The expansion of pericorneal stem cells from a proximal lineage (black trace) compensates for the cell loss and resume the centripetal regeneration of that segment of the cornea. Arrows indicate the site of ablation. (**F)** High magnification views of the same ablation site showing no contribution from cells in the outer limbus. Asterisk marks the site of ablation and dotted lines indicate the border between the outer and pericorneal limbus. (**G)** Proposed model of the bi-compartmentalized stem cell organization of the limbal niche based on the observed stem cell behaviors and their respective contributions to tissue regeneration. Scale bars: 200 μm.

We next asked which cells compensate for the loss of corneal stem cells following ablation. In these experiments, we consistently captured stem cells from separate proximal corneal lineages entering the ablation site and commence centripetal growth, increasing their own clone size to compensate (**Fig. 4E)**. In contrast, cells from the smaller clones in the outer limbus did not cross over into the pericorneal limbus or contribute to the long-term regeneration of the cornea **(Fig. 4F)**. Taken together, our results show that the label-retaining cells found in the outer limbus are functionally distinct from the corneal stem cells and may play a role in maintaining a border between the cornea and the conjunctiva **(Fig. 4G)**.

In this study, we resolve the long-standing question of the functional corneal stem cell reservoir. Our data reveal the bi-compartmentalized organization of the limbus and support a model in which corneal stem cells are located in a distinct niche, the pericorneal limbus **(Fig. 3D; 4G)**. Stem cells in the pericorneal limbus are necessary and sufficient to regenerate the cornea, and are the sole source of the short-lived basal progenitors found in the rest of the tissue. Furthermore, slow-cycling cells are specifically located in the outer limbus. These cells do not contribute to the cornea, but form distinct lineages oriented circumferentially along the limbus. This behavior suggests that slow-cycling cells in the outer limbus may be required to maintain an impenetrable barrier from the conjunctiva. Such functionally distinct stem cell populations appear to be common in transition areas between epithelia in other organs (*34*). Our study provides a paradigm for the use of live imaging to elucidate fundamental mechanisms of epithelial biology and to uncover how compartmentalized stem cell activity supports tissue function and regeneration.

## Acknowledgments

We thank Michael Rendl, Christopher Lengner and Kenneth Zaret for our invaluable discussions and their critical comments on the manuscript. We are especially grateful to George Cotsarelis for many helpful conversations that guided this study. We thank Elaine Fuchs for donating the *K14*^*H2B-GFP*^ mouse line and the *K14*-cassette vector.

## Funding

O.F. was supported by training grant T32HD083185 from NIH/NICHD. Y.S.H. was supported by training grant T32AR00746536 from NIH/NIAMS. G.R. was supported by training grant T32GM007229 from NIH/NIGMS. P.K. was supported by the American Association for Cancer Research-John and Elizabeth Leonard Family Foundation Basic Cancer Research Fellowship. V.L. was supported by a grant from NIH/NEI (K08 EY025742). P.R. was supported by grants from NIH/NEI (R01EY030599) and from the American Cancer Society (RSG1803101DCC).

## Author contributions

O.F. and P.R. conceptualized the overall study, designed the experiments and wrote the manuscript. O.F., Y.H.S., M.B., P.K., S.H., G.R. and P.R. performed the experiments. J.X. provided the *p63*^*CreER*^ mouse line, T.D. provided technical support with histology, V.L. provided human cornea samples and technical support. All authors discussed results and participated in the manuscript preparation and editing. P.R. supervised the study.

## Competing interests

The authors declare no competing interests.

## Data and materials availability

Data sets and reagents presented in this study are available from the corresponding author upon request.

## Supplementary Materials

### Materials and Methods

#### Mice

All procedures involving animal subjects were performed with the approval of the Institutional Animal Care and Use Committee (IACUC) of the University of Pennsylvania and were consistent with the guidelines set forth by the AVRO Statement for the Use of Animals in Ophthalmic and Vision Research. *K14*^*H2B-GFP*^ mice were generated by E. Fuchs (*19*). *p63*^*CreER*^ mice were created by Jianming Xu (*36*). *R26*^*loxp-stop-loxp-tdTom*^ (Stock No #007908), *R26*^*loxp-mTom-stop-loxp-mGFP*^ (*R26*^*mTmG*^ in the text; Stock No #007676) and *R26*^*loxp-nTom-stop-loxp-nGFP*^ (*R26*^*nTnG*^ in the text; Stock No #023035) mice were obtained from Jackson Laboratories. *R26*^*PAGFP*^ reporter mice were generated by crossing *E2a*^*Cre*^ (Stock No #003724) with *R26*^*loxp-stop-loxp-PAGFP*^ (Stock No #021071), to achieve germline transmission of the recombined allele and ubiquitous expression of the PAGFP reporter. Cre activation was induced with a single intraperitoneal injection of Tamoxifen in corn oil (0.2-2 mg per 20 g body weight). Mice between 2-6 months old were used for experiments, with equal representation of males and females. Mice were housed under standard laboratory conditions and received food and water *ad libitum*.

#### Generation of *K14*^*H2B-PAGFP*^ mice

An H2BPAGFP coding sequence was obtained from the pACAGW-H2B-PAGFP-AAV plasmid (M.Scanziani via Addgene, #33000) by PCR amplification and subcloned into the pG3Z-K14-H2B vector (E. Fuchs) by restriction enzyme digestion with BamHI and XbaI (NEB) and ligation with T4 Ligase (NEB). The final K14-H2BPAGFP transgene was obtained from the resultant plasmid by digestion with KpnI and SphI and injected into blastocysts by the Center for Animal Transgenesis and Germ Cell Research, at the School of Veterinary Medicine of the University of Pennsylvania. Resulting mice were first screened by whole mount imaging and photo-activation of a freshly obtained ear punch biopsy, using our 2-photon microscope and the positive genotypes were confirmed using specific primers for the *K14* cassette. A single founder was then selected and crossed with a CD-1 mixed background breeder (Charles River, Stock No #022) to establish the transgenic mouse line.

#### Intravital imaging of the mouse eye

Preparation of the mice for intravital imaging of the eye was performed with the following amendments to the previously described protocol (*37*). Mice were initially anesthetized with IP injection of ketamine/xylazine cocktail (0.1 ml / 20 g body weight; 87.5 mg / kg Ketamine, 12.5 mg / kg Xylazine). A deep plane of anesthesia was verified by checking pedal reflexes. The mouse head was stabilized with a custom-made stereotaxic apparatus that includes palate bar and nose clamp but no ear bars. Precision, 3-axis micro-manipulators are used to adjust the head tilt so that the eye to be imaged is facing up. A drop of eye gel (0.3 % Hypromellose) was used as an optically neutral interface between the eye and a glass coverslip, and to prevent dryness and irritation to the tissue during the anesthesia and imaging procedure. After preparation and mounting is complete, the stage is placed on the microscope platform under the objective lens. A heating pad is used to keep a stable body temperature and vaporized isoflurane is delivered through a nose cone to maintain anesthesia for the duration of the imaging process. After each imaging session, the eyes were rinsed with PBS and the mice were monitored and allowed to recover in a warm chamber before returned to the housing facility.

#### Imaging Equipment and Acquisition Settings

Image acquisition was performed with an upright Olympus FV1200MPE microscope, equipped with a Chameleon Vision II Ti:Sapphire laser. The laser beam was focused through 10X, 20X or 25X objective lenses (Olympus UPLSAPO10×2, N.A. 0.40; UPLSAPO20X, N.A. 0.75; XLPLN25XWMP2, N.A. 1.05) and. Emitted fluorescence was collected by two multi-alkali and two gallium arsenide phosphide (GaAsP) non-descanned detectors (NDD). The following wavelengths were collected by each detector: NDD1 419-458 nm; NDD1 458–495 nm; GaAsP-NDD1 495–540 nm; GaAsP -NDD2 575–630 nm. GFP and Tomato reporters were excited at 930 nm and their signal was collected by GaAsP-NDD1 and GaAsP-NDD2, respectively. Second harmonic generation (SHG) signal was generated using 850nm or 930 nm excitation wavelengths and detected in NDD1 or NDD2, respectively. Serial optical sections were acquired in 2–5 μm steps, starting from the surface of the eye and capturing the entire thickness of the cornea (epithelium ∼40 μm, stroma/endothelium ∼80 μm). Multi-day tracing experiments were done by re-imaging the same field of view or the entire eye at the indicated times after the initial acquisition. For each time point, inherent landmarks within the cornea including the organization of the vasculature and collagen fibers (SHG) were used to consistently identify the limbus and navigate back to the original regions.

#### Photo-Labeling

Photo-labeling experiments with the *K14*^*H2BPAGFP*^ and *R26*^*PAGFP*^ reporter mice were carried out with the same equipment and imaging setup as used for acquisition. The pre-activated form of the H2B-PAGFP and PAGFP fluorescent proteins was visualized by exciting with 850nm wavelength and emission signal was collected in GaAsP-NDD1. Excitation with 930nm verified that no signal is emitted by the reporters before activation. Photo-labeling was achieved by scanning a defined region-of-interest (ROI) at the plane of the basal layer of the epithelium, with the laser tuned to 750nm wavelength, for 5-10 sec, using 5-10% laser power. The z-plane was then moved down to the corneal endothelium, which served as a reference, and the same ROI was used for photo-labeling cells in that layer. Immediately after photo-activation, a series of optical sections, with a range that includes the entire thickness of the cornea, were acquired using the same acquisition settings as for GFP. Visualizing the signal of the activated form of PAGFP only within the ROI confirmed the successful photo-labeling of basal epithelial or endothelial cells. Following the initial image acquisition immediately after photo-labeling the same eyes were re-imaged at the indicated times to evaluate the changes of the labeled epithelial population and their movements compared to the endothelial reference cell group.

#### Laser Ablation

*In vivo* laser-induced cell ablation was performed using the same femtosecond Ti:Sapphire laser used for fluorescence excitation and imaging. For maximum specificity, a 25X objective lens (XLPLN25XWMP2, N.A. 1.05) was used to focus the laser beam to the basal layer of the eye epithelium. The laser was guided to scan a small area (∼100 μm^2^), targeting the cells within for ablation with the following parameters: 800nm wavelength, 100% laser power, 1s exposure. Immediately after ablation, the microscope was switched to imaging mode and a series of serial optical sections were collected to visualize the effect of the laser ablation.

#### Quantitative Image Analysis

Raw digital files from 2-photon imaging were acquired and saved in the OIB format using the microscope manufacturer’s software (FLUOVIEW, Olympus USA). To capture extended fields-of-view that encompass the entire eye surface epithelium, including the cornea, limbus and conjunctiva a tiling method was used to reconstruct a single image from multiple full-thickness serial optical sections, using the microscope acquisition software. Typically, to image the entire eye using the 10X objective lens the microscope defines a square area consisting of 2 × 2 (XY) fields-of-view, with 10% overlap between them. Using the motorized platform, the microscope automatically acquires the four fields-of-view in a sequential pattern and uses information from the overlapping margins to stitch the individual fields-of-view into a single image. Raw image files were imported into ImageJ/Fiji (NIH) using Bio-Formats to convert them to TIFF files and for further analysis. For cell counts and quantitative clonal analyses, supervised image segmentation and blob detection was performed on individual optical sections. identified blobs were manually validated and their number, size and signal intensity, as mean grey values were measured. Clonal growth anisotropy was analyzed by measuring Feret’s Diameter and Angles, from the outlines of individual clones after image segmentation. Clone measurements and tracings for the cell ablation experiments were performed manually. Images shown in figures typically represent maximum projections or single optical sections selected from the z-stacks, unless otherwise specified.

#### Histology and immunolabeling

Mouse eyes were fixed in 4% paraformaldehyde overnight at 4°C, then placed in 30% sucrose for 60 min. Eyes were embedded and frozen in Tissue-Tek OCT Compound (Sakura Finetek). 10 μm sections were stained with Keratin 14 (Biolegend, rabbit 1:1000) and DAPI (Biotium, 1:5000). Human corneas were fixed in 10% formalin overnight at 4°C, then cut into six or eight wedges. Each piece was permeabilized with 0.3% Triton X-100 and stained with Phalloidin (Biotium, 1:1000) and DAPI (Biotium, 1:1000). Human corneal sections were retrieved from the archives of the William C. Frayer Ocular Pathology Laboratory, Department of Ophthalmology, at the University of Pennsylvania.

#### Statistical analysis

Sample sizes were not pre-determined, but are similar with what was reported previously (*5*). Data were collected and quantified randomly and their distribution was assumed normal, but this was not formally tested. Lineage tracing, photo-labeling and laser-ablation experiments were performed and successfully reproduced at least three times using different mice. The values of “N” (sample size) refer to data points from a single experiment, unless otherwise indicated, and are provided in the figure legends. Statistical calculations and graphical representation of the data were performed using the Prism software package (GraphPad). Data are expressed as percentages or mean ± S.E.M and unpaired Student’s *t*-test was used to analyze data sets with two groups, unless otherwise stated in the figure legends. For all analyses, *P*-values < 0.05 were designated as significant and symbolized in figure plots as \**P* < 0.05, \*\**P* < 0.01, \*\*\**P* < 0.001, \*\*\*\**P* < 0.0001, with precise values supplied in figure legends. No data were excluded from the analysis.

**Figure S1.**
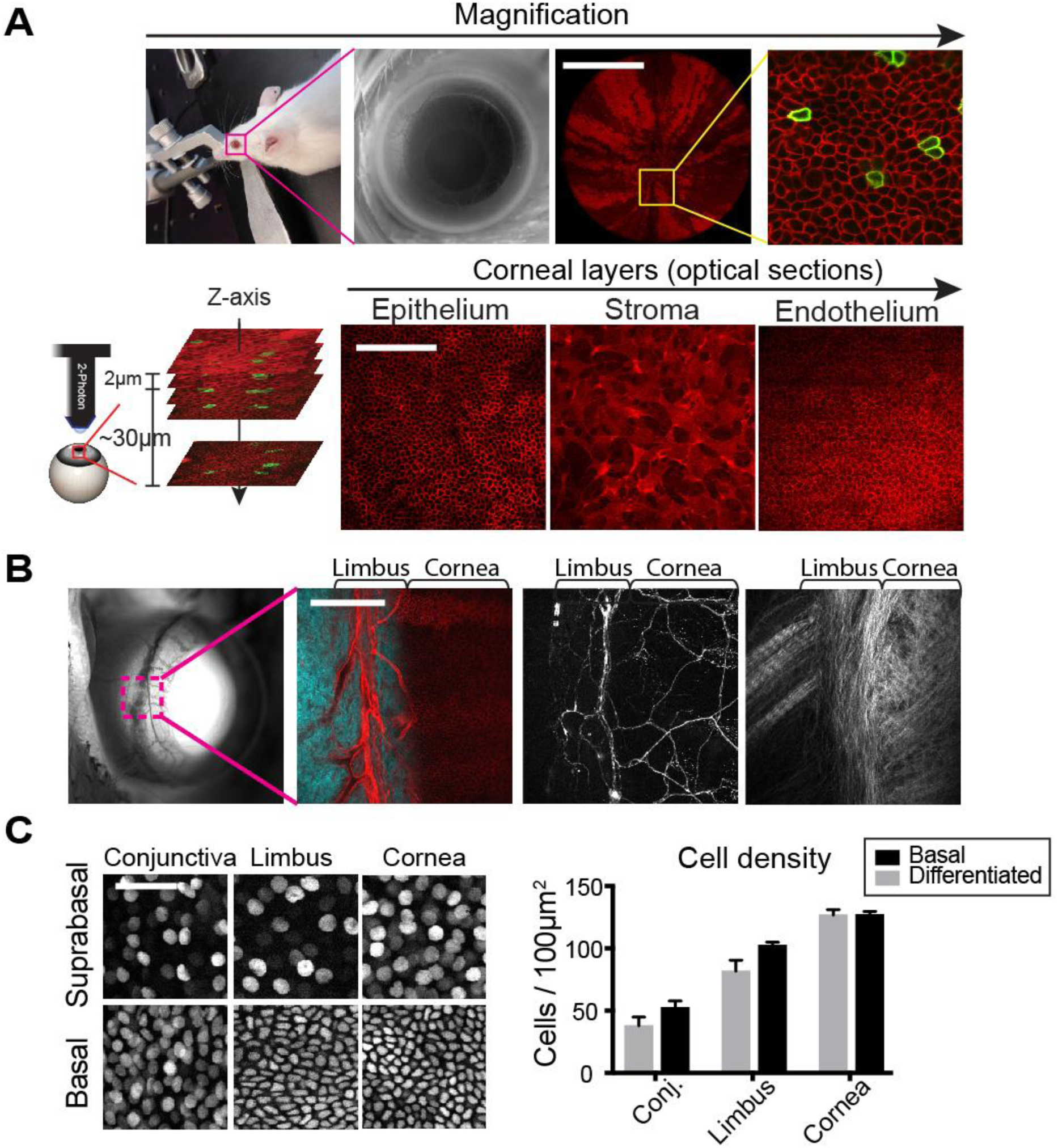
*In vivo* 2-photon imaging of the mouse eye. **(A)** Increasing magnification of our imaging approach (upper panel). Scale bar: 1mm. Individual planes of the corresponding tissue compartments, taken from serial optical sections of the live cornea (lower panel). Scale bar: 200µm. **(B)** Images demonstrate co-localization of blood vessels (left), sensory nerves (middle), and extracellular matrix (right), within the limbal niche. Scale bar: 200µm. **(C)** Images and corresponding quantification of the total number of basal and suprabasal cells compared across the eye surface epithelium. Scale bar: 50 µm.

**Figure S2.**
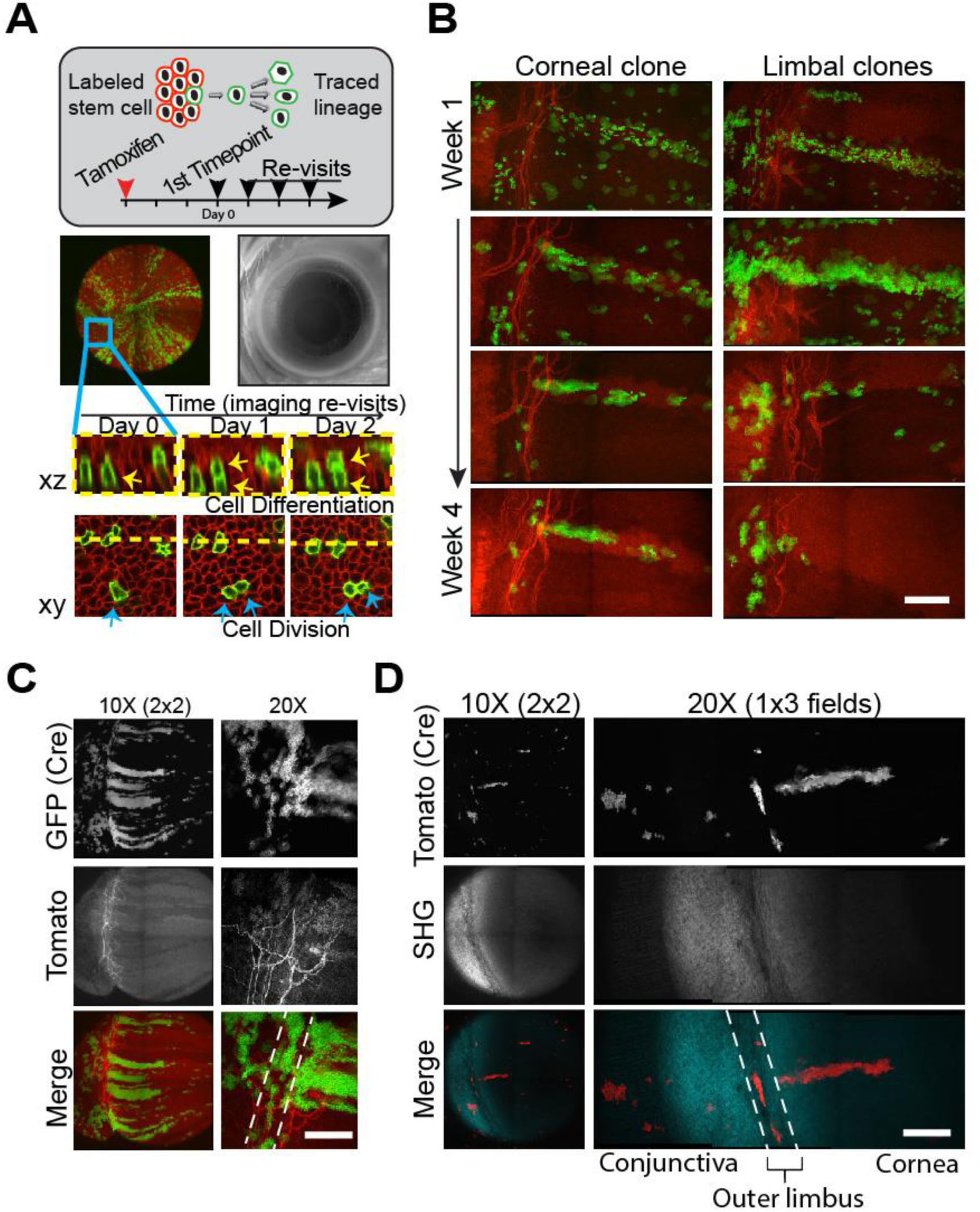
Visualizing distinct clonal dynamics by in vivo lineage tracing and live imaging. **(A)** Lineage tracing and imaging strategy to track long-term activity of single cells in the cornea. **(B)** A single radially oriented corneal clone that persists through week 4 post-induction (left panel). Smaller limbal clones that remain in place and do not contribute to the regeneration of the cornea (right panel). **(C, D)** Representative examples of distinct clonal growth patterns in the cornea, limbus and conjunctiva. Dotted lines mark the location of the outer limbus. Scale bars: 200 µm.

**Figure S3.**
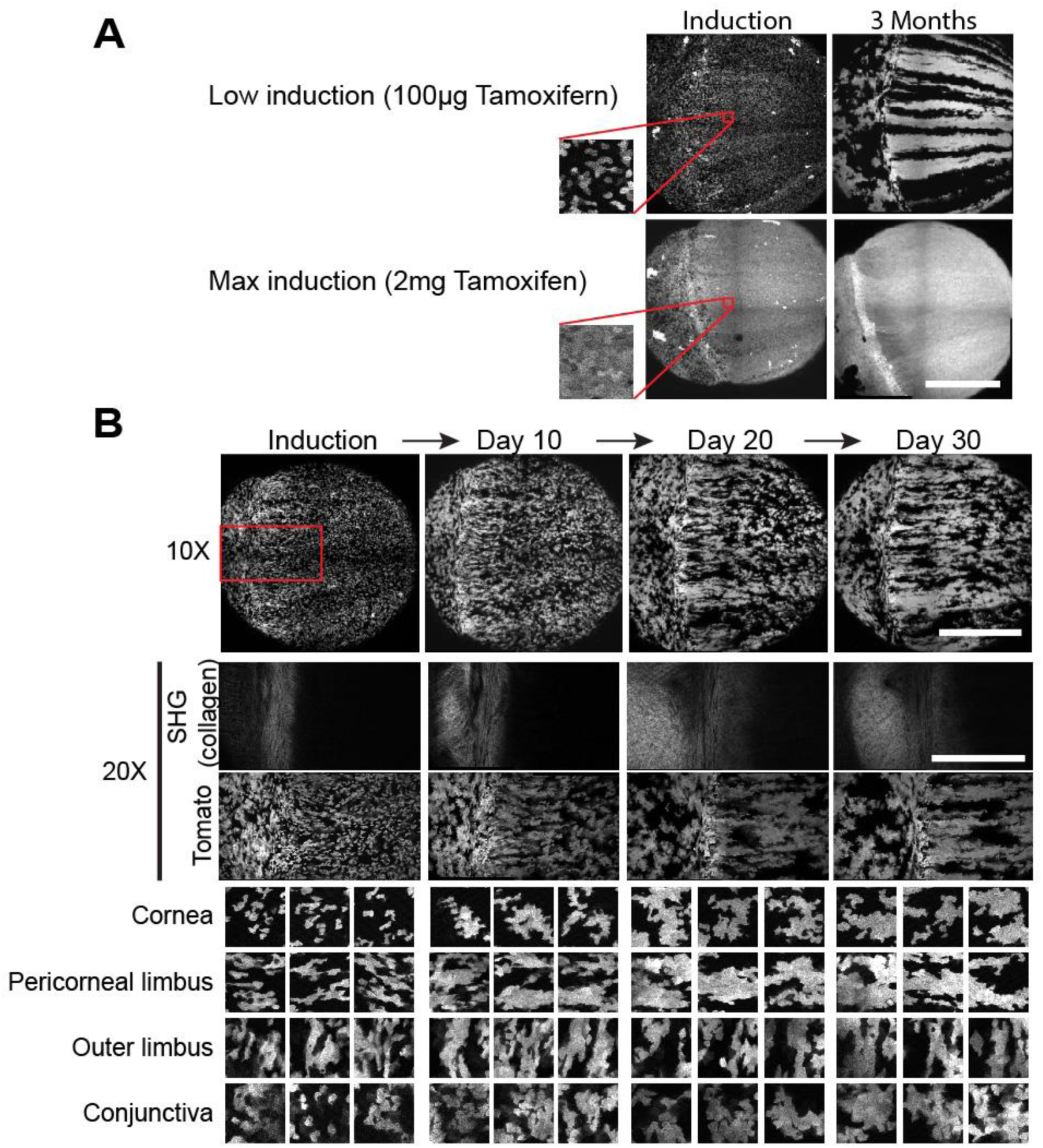
Live imaging of epithelial clonal dynamics. **(A)** Representative images of the eyes of p63^CreER;^ R26^loxp-stop-loxp-tdTom^ mice after low or max Tamoxifen induction. Scale bar: 1 mm. **(B)** Images of the whole eye surface epithelium every 10 days during a 30-day lineage tracing time course (top panel). Scale bar: 1mm. Area outlined in red in higher magnification (middle panels). Scale bar: 500 µm. Examples of distinct clonal patterns visualized in different areas of the eye surface epithelium (lower panels).

**Figure S4.**
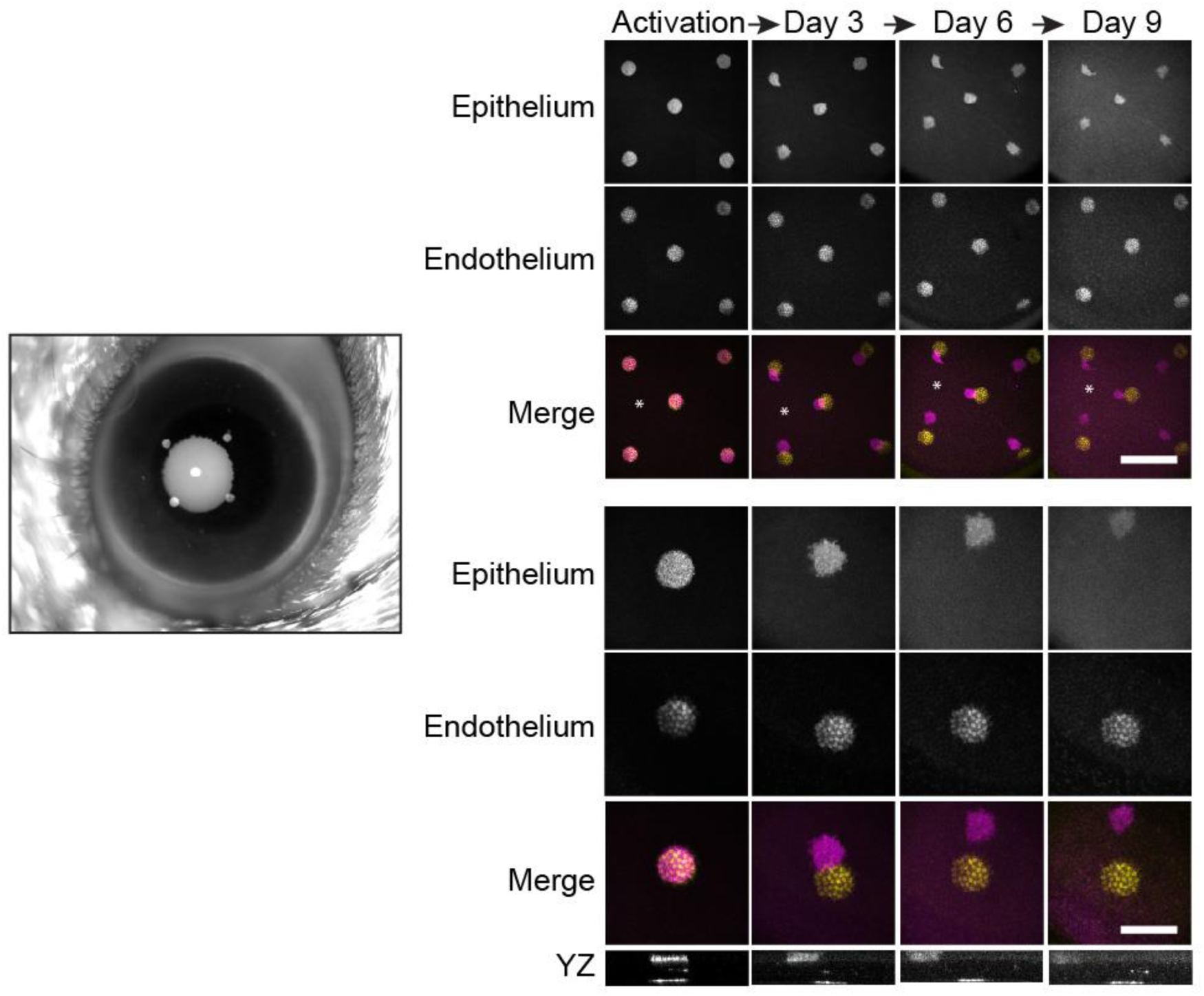
Direct capture of the centripetal movement of corneal progenitors. Five groups of basal epithelial cells within the central cornea were unbiasedly photo-labeled, using a globally expressed photo-activatable, GFP reporter. Endothelial cells immediately below were used as a reference. Top right panels show low magnification view of the central cornea where the photo-labeled groups of cells can be seen during the time course. The movement of the labeled cells was visualized every 3 days over the course of 9 days. Basal cells in the central cornea moved centripetally in unison towards the center of the eye (asterisk). Scale bar: 500 µm. Bottom right panels show one example of a group of labelled epithelial and endothelial cells. Scale bar: 200 µm.

**Figure S5.**
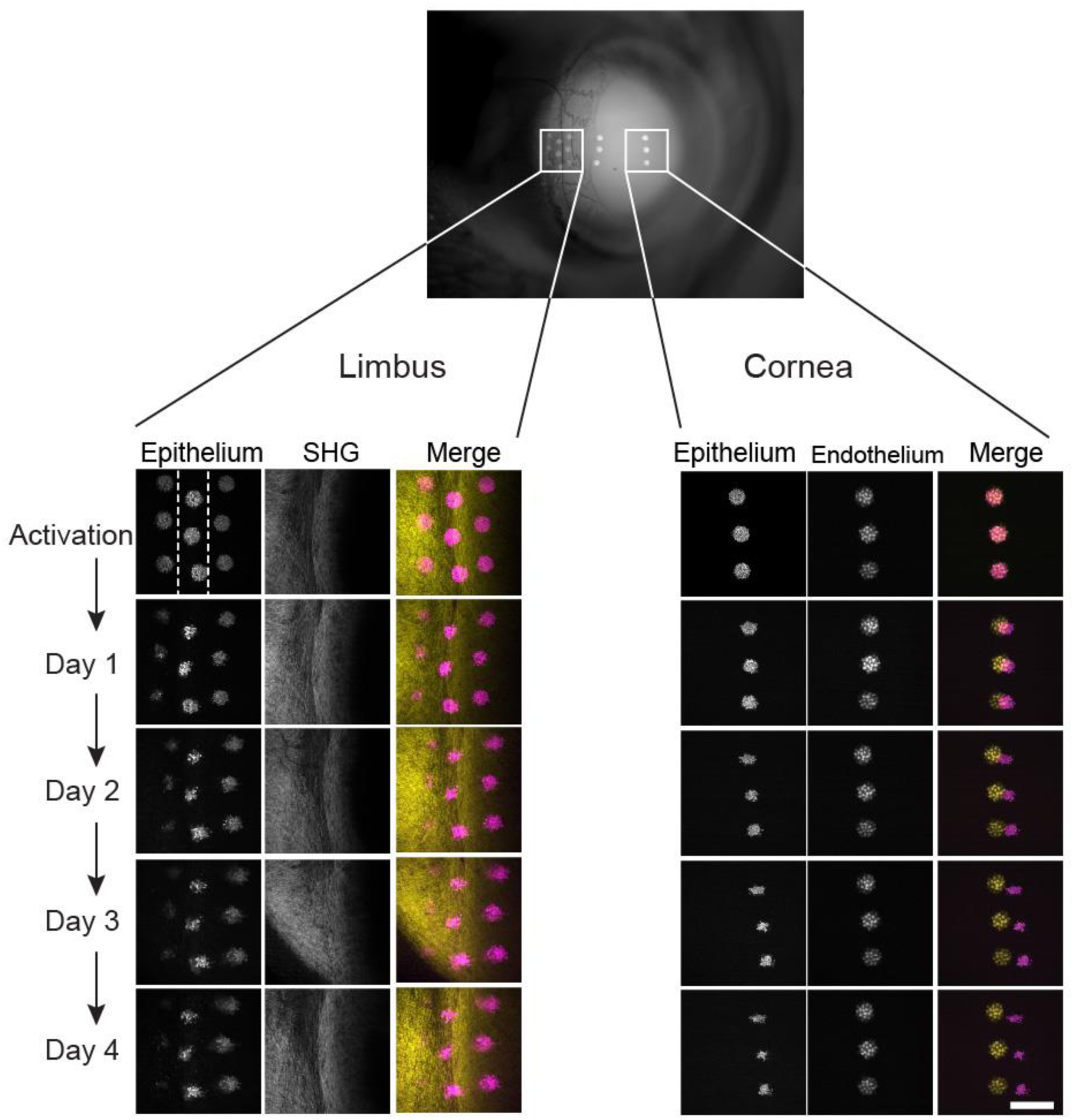
Capturing dynamic cellular mobility in the regenerating cornea. Time course images of labeled cells in the epithelium and endothelium of the central cornea (left panels). By 4 days after labeling, epithelial cells have moved away from their original location towards the center of the eye. Groups of labeled epithelial cells in the pericorneal and outer limbus as well as in the conjunctiva (right panels). SHG imaging is used to visualize the distinctive limbal collagen fibers in the underlying stroma, and is used as a reference. Pericorneal cells begin to move away from the limbus and towards the center of the eye. Limbal cells remain in their miginal location. Conjunctival cells also do not move, but undergo rapid turnover, so that the GFP signal is barely visible after 4 days of tracing. Scale bar: 200 µm.

**Figure S6.**
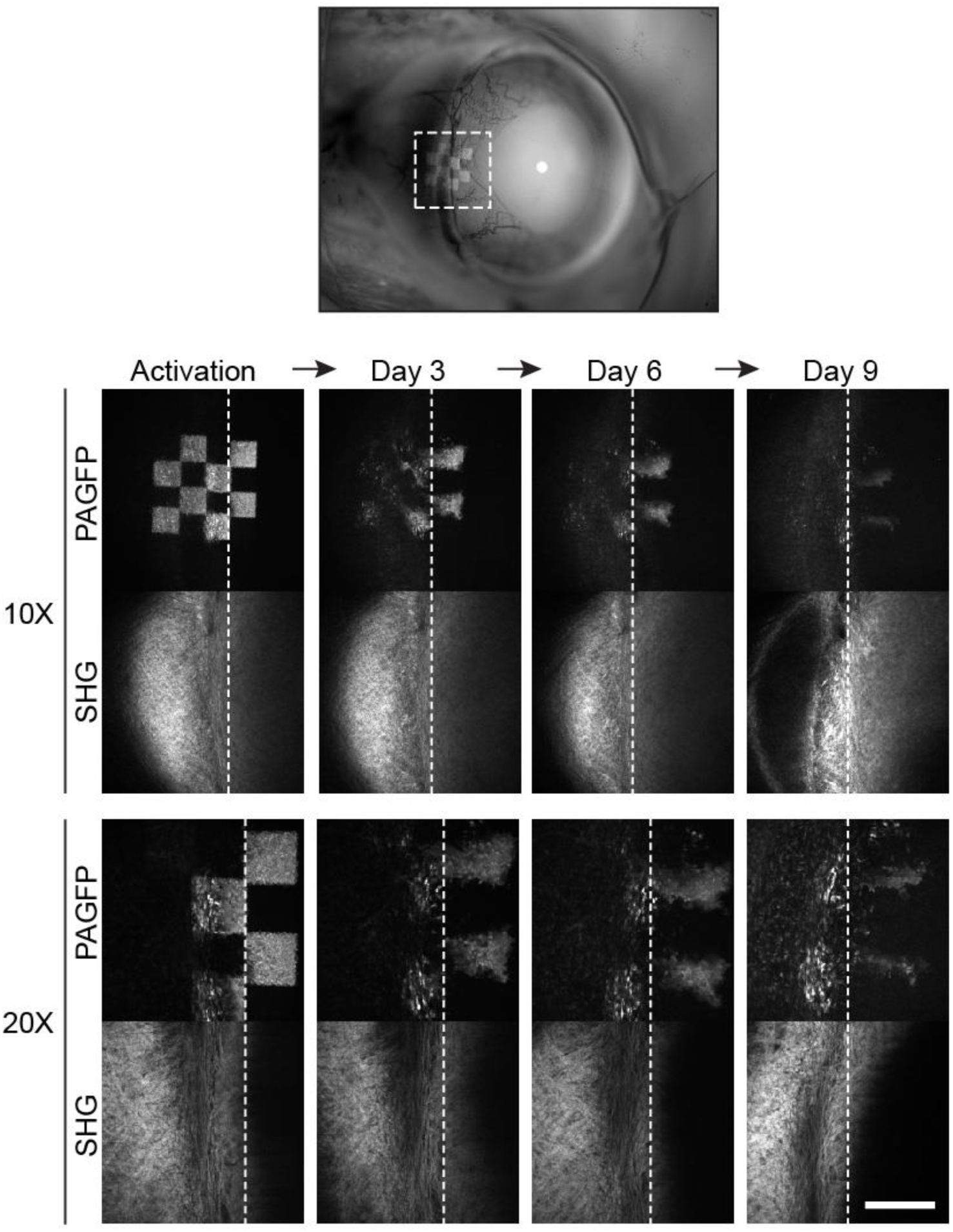
Tracking cellular dynamics of limbal cells. All images from the full experimental time course shown in Fig. 2C. Changes in the checkered pattern shows centripetal expansion in the pericomeal limbus (dotted line), but not the outer limbus or conjunctiva. SHG imaging is used to visualize the distinctive limbal collagen fibers in the underlying stroma, and is used as a reference. Scale bar: 200 µm.

**Figure S7.**
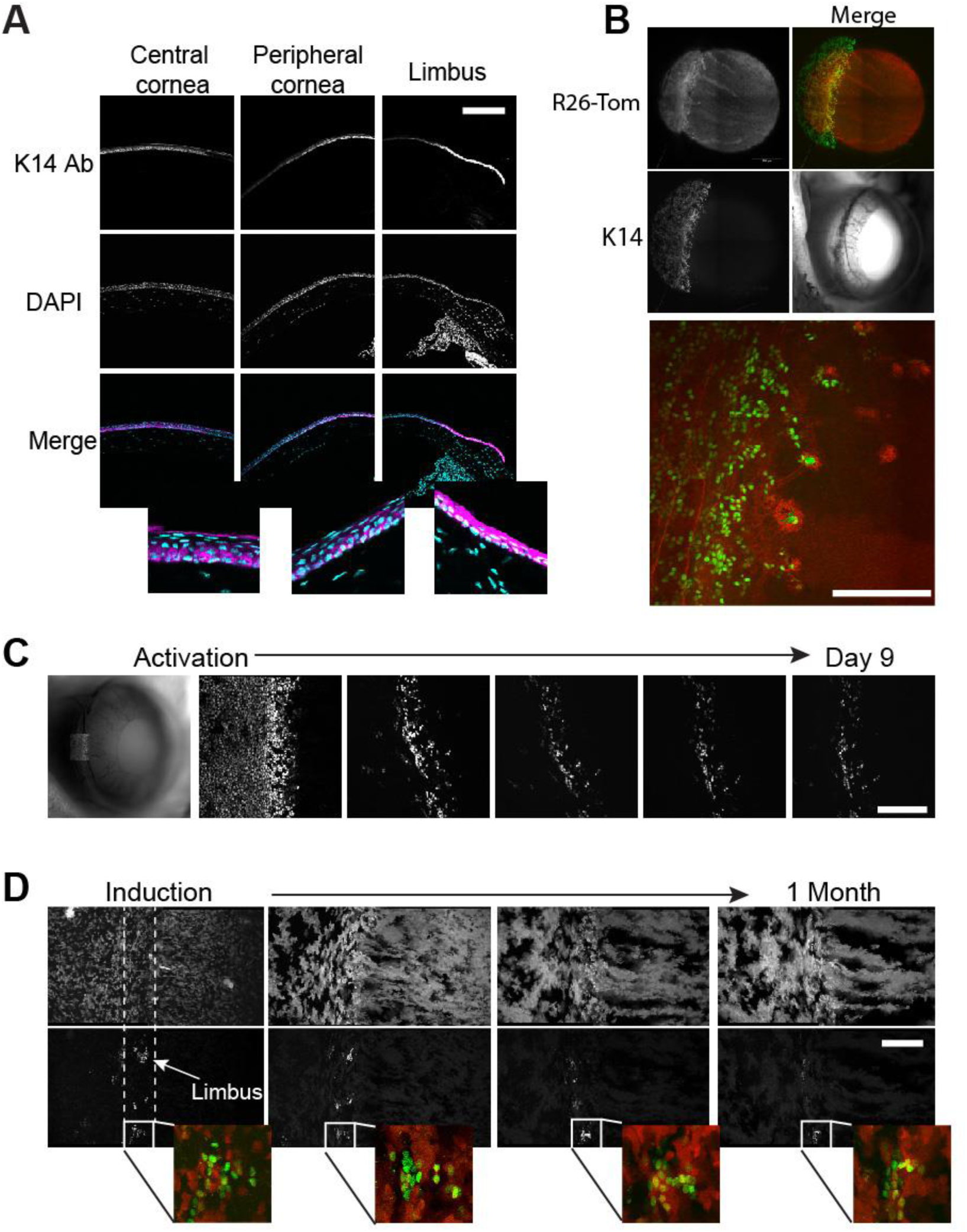
Visualizing label retaining cells in the live eye. **(A)** Imnmnostained sections showing the compartmentalized high expression of Keratin 14 in the limbus compared to the central cornea. **(B)** Low and high magnification images of the live eye from a transgenic Kl4H2B-GFP reporter mouse line showing similar high expression in the limbal area. **(D, E)** All timepoints from the experiments shown in Fig. 3C and 3D, respectively. Scale bars: 200 µm.

**Figure S8.**
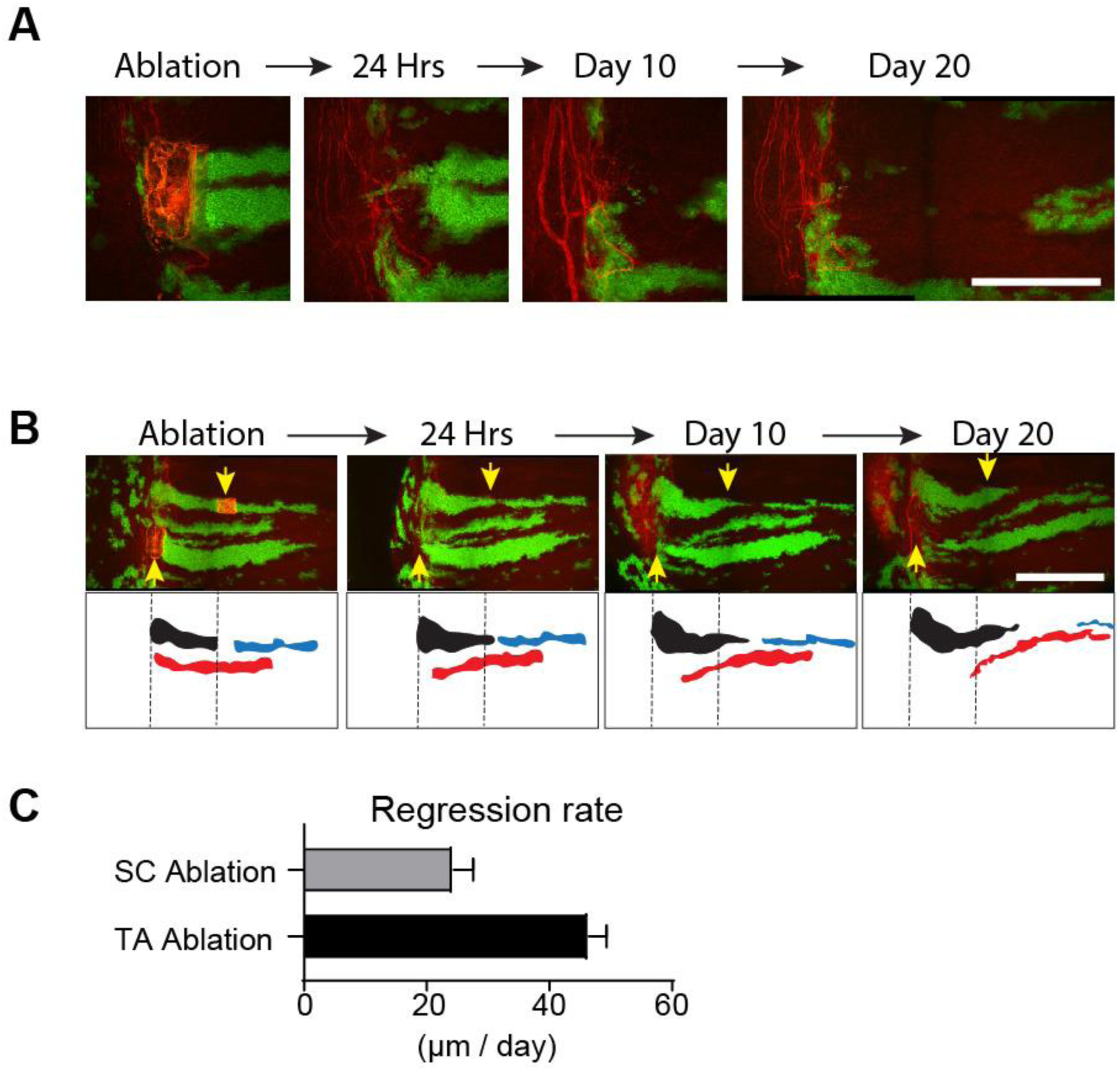
2-photon ablation and lineage tracing of corneal stem cells and progenitors. **(A)** High magnification images from the experimental time course shown in Fig. 4C. **(B)** Injury does not reverse the centripetal movement and fate of basal corneal progenitors. 2-photon images of the limbus and cornea taken at the indicated time points after cell ablation (top panel). Traces indicate segments of a corneal lineage upstream (black) or down-stream (blue) of the of the ablated progenitor cells. A separa te lineage (red) with ablated stem cells in the pericorneal limbus is also shown for comparison. Arrows indicate the site of ablations. **(C)** Quantification of the centripetal regression rate of corneal lineages down-stream of the stem cells and progenitors ablation sites. Scale bars: 500µm.

**<u>Movie S1</u>. Intravital 2-photon imaging of the mouse eye**. Example of a mouse eye, imaged live by 2-photon microscopy. Serial optical sections are obtained in sequential fields of view and the entire anterior segment of the eye can be reconstructed in two or three dimensions at single cell resolution.

**<u>Movie S2</u>. Characterization of cellular morphology in the eye by live imaging**. Example of serial optical sections of the conjunctiva, limbus and cornea, imaged live using the *R26*^*mTmG*^ reporter mouse line, which illuminates the membrane of all cells in the tissue.

**<u>Movie S3</u>. Characterization of cellular number and density in the eye by live imaging**. Example of serial optical sections of the conjunctiva, limbus and cornea, imaged live using the *R26*^*nTnG*^ reporter mouse line, which illuminates the nucleus of all cells in the tissue.

**<u>Movie S4</u>. Characterization of extracellular matrix organization in the eye by live imaging**. Example of serial optical sections of the conjunctiva, limbus and cornea, imaged live using the second harmonic generation imaging (SHG), which illuminates the collagen fibers in the tissue.

**<u>Movie S5</u>. Conserved organization of the mammalian limbus**. Example of serial optical sections of a whole-mount human limbus, stained with 1) phalloidin, which marks cortical actin and illuminates the outlines of all cells, 2) DAPI, which marks the nuclei of all cells and 3) second harmonic generation imaging (SHG) which marks the collagen in the stroma of the tissue.

**<u>Movie S6</u>. Visualizing and tracking dynamic cellular behaviors by *in vivo* photo-labeling**. Example of a group of corneal basal progenitors marked by photo-activating the *R26*^*PAGFP*^ reporter by 2-photon microscopy. Serial optical sections immediately after labeling show the group of epithelial cells located directly above a similarly sized group of corneal endothelial cells which are quiescent and are used as control. Four days later the group of epithelial cells has moved towards the center of the cornea.

## Notes

### Competing Interest Statement

The authors have declared no competing interest.

### Summary of Updates

Authors list

## References and Notes

1. Marques-pereira JP, Leblond CP. Mitosis and differentiation in the stratified squamous epithelium of the rat esophagus. Am J Anat. 1965;117:73–87.

2. Green H. The keratinocyte as differentiated cell type. Harvey Lect. 1980;74:101–39.

3. Clayton E, Doupé DP, Klein AM, Winton DJ, Simons BD, Jones PH. A single type of progenitor cell maintains normal epidermis. Nature. 2007;446(7132):185–9.

4. Mascré G, Dekoninck S, Drogat B, et al. Distinct contribution of stem and progenitor cells to epidermal maintenance. Nature. 2012;489(7415):257–62.

5. Rompolas P, Mesa KR, Kawaguchi K, et al. Spatiotemporal coordination of stem cell commitment during epidermal homeostasis. Science. 2016;352(6292):1471–4.

6. Schermer A, Galvin S, Sun TT. Differentiation-related expression of a major 64K corneal keratin in vivo and in culture suggests limbal location of corneal epithelial stem cells. J Cell Biol. 1986;103(1):49–62.

7. Cotsarelis G, Cheng SZ, Dong G, Sun TT, Lavker RM. Existence of slow-cycling limbal epithelial basal cells that can be preferentially stimulated to proliferate: implications on epithelial stem cells. Cell. 1989;57(2):201–9.

8. Pellegrini G, Golisano O, Paterna P, et al. Location and clonal analysis of stem cells and their differentiated progeny in the human ocular surface. J Cell Biol. 1999;145(4):769–82.

9. Notara M, Alatza A, Gilfillan J, et al. In sickness and in health: Corneal epithelial stem cell biology, pathology and therapy. Exp Eye Res. 2010;90(2):188–95.

10. Scadden DT. Nice neighborhood: emerging concepts of the stem cell niche. Cell. 2014;157(1):41–50.

11. Huang AJ, Tseng SC. Corneal epithelial wound healing in the absence of limbal epithelium. Invest Ophthalmol Vis Sci. 1991;32(1):96–105.

12. Lavker RM, Dong G, Cheng SZ, Kudoh K, Cotsarelis G, Sun TT. Relative proliferative rates of limbal and corneal epithelia. Implications of corneal epithelial migration, circadian rhythm, and suprabasally located DNA-synthesizing keratinocytes. Invest Ophthalmol Vis Sci. 1991;32(6):1864–75.

13. Collinson JM, Morris L, Reid AI, et al. Clonal analysis of patterns of growth, stem cell activity, and cell movement during the development and maintenance of the murine corneal epithelium. Dev Dyn. 2002;224(4):432–40.

14. P. Rama, S. Matuska, G. Paganoni, A. Spinelli, M. Luca, G. Pellegrini, Limbal stem-cell therapy and long-term corneal regeneration. New Engl J Medicine. 363, 147–155 (2010).

15. Di girolamo N, Bobba S, Raviraj V, et al. Tracing the fate of limbal epithelial progenitor cells in the murine cornea. Stem Cells. 2015;33(1):157–69.

16. Amitai-lange A, Altshuler A, Bubley J, Dbayat N, Tiosano B, Shalom-feuerstein R. Lineage tracing of stem and progenitor cells of the murine corneal epithelium. Stem Cells. 2015;33(1):230–9.

17. N. J. Dorà, R. E. Hill, M. J. Collinson, J. D. West, Lineage tracing in the adult mouse corneal epithelium supports the limbal epithelial stem cell hypothesis with intermittent periods of stem cell quiescence. Stem Cell Res. 15, 665–677 (2015).

18. Cotsarelis G, Sun TT, Lavker RM. Label-retaining cells reside in the bulge area of pilosebaceous unit: implications for follicular stem cells, hair cycle, and skin carcinogenesis. Cell. 1990;61(7):1329–37.

19. Tumbar T, Guasch G, Greco V, et al. Defining the epithelial stem cell niche in skin. Science. 2004;303(5656):359–63.

20. A. Wilson, E. Laurenti, G. Oser, R. C. van der Wath, W. Blanco-Bose, M. Jaworski, S. Offner, C. F. Dunant, L. Eshkind, E. Bockamp, P. Lió, R. H. MacDonald, A. Trumpp, Hematopoietic stem cells reversibly switch from dormancy to self-renewal during homeostasis and repair. Cell. 135, 1118–1129 (2008).

21. Takeda N, Jain R, Leboeuf MR, Wang Q, Lu MM, Epstein JA. Interconversion between intestinal stem cell populations in distinct niches. Science. 2011;334(6061):1420–4.

22. P. Rompolas, E. R. Deschene, G. Zito, D. G. Gonzalez, I. Saotome, A. M. Haberman, V. Greco, Live imaging of stem cell and progeny behaviour in physiological hair-follicle regeneration. Nature. 487, 496 (2012).

23. P. Rompolas, K. R. Mesa, V. Greco, Spatial organization within a niche as a determinant of stem-cell fate. Nature. 502, 513 (2013).

24. S. Huang, P. Rompolas, Two-photon microscopy for intracutaneous imaging of stem cell activity in mice. Exp Dermatol. 26, 379–383 (2017).

25. Van buskirk EM. The anatomy of the limbus. Eye (Lond). 1989;3 (Pt 2):101–8.

26. X. Chen, O. Nadiarynkh, S. Plotnikov, P. J. Campagnola, Second harmonic generation microscopy for quantitative analysis of collagen fibrillar structure. Nat Protoc. 7, 654 (2012).

27. Doupé DP, Alcolea MP, Roshan A, et al. A single progenitor population switches behavior to maintain and repair esophageal epithelium. Science. 2012;337(6098):1091–3.

28. Allen TD, Potten CS. Fine-structural identification and organization of the epidermal proliferative unit. J Cell Sci. 1974;15(2):291–319.

29. Majo F, Rochat A, Nicolas M, Jaoudé GA, Barrandon Y. Oligopotent stem cells are distributed throughout the mammalian ocular surface. Nature. 2008;456(7219):250–4.

30. West JD, Dorà NJ, Collinson JM. Evaluating alternative stem cell hypotheses for adult corneal epithelial maintenance. World J Stem Cells. 2015;7(2):281–99.

31. R. Sartaj, C. Zhang, P. Wan, Z. Pasha, V. Guaiquil, A. Liu, J. Liu, Y. Luo, E. Fuchs, M. Rosenblatt, Characterization of slow cycling corneal limbal epithelial cells identifies putative stem cell markers. Sci Rep. 7, 3793 (2017).

32. A. Richardson, E. P. Lobo, N. C. Delic, M. R. Myerscough, G. J. Lyons, D. Wakefield, N. Girolamo, Keratin-14-positive precursor cells spawn a population of migratory corneal epithelia that maintain tissue mass throughout life. Stem Cell Rep. 9, 1081–1096 (2017).

33. Pajoohesh-ganji A, Pal-ghosh S, Tadvalkar G, Stepp MA. K14 + compound niches are present on the mouse cornea early after birth and expand after debridement wounds. Dev Dyn. 2016;245(2):132–43.

34. Runck LA, Kramer M, Ciraolo G, Lewis AG, Guasch G. Identification of epithelial label-retaining cells at the transition between the anal canal and the rectum in mice. Cell Cycle. 2010;9(15):3039–45.

## Methods References

35. Lee DK, Liu Y, Liao L, Wang F, Xu J. The prostate basal cell (BC) heterogeneity and the p63-positive BC differentiation spectrum in mice. Int J Biol Sci. 2014;10(9):1007–17.

36. Pineda CM, Park S, Mesa KR, et al. Intravital imaging of hair follicle regeneration in the mouse. Nat Protoc. 2015;10(7):1116–30.

